# *Staphylococcus aureus* adopts revertible intracellular viable-but-non-culturable phenotypes in chronic osteocyte infections

**DOI:** 10.1101/2024.12.17.629047

**Authors:** Nicholas J. Gunn, Anja R. Zelmer, M. Amjad Hossain, Kate P. Whyte, L. Bogdan Solomon, Miguel Carda-Diéguez, Alex Mira, Anna E. Sheppard, Stephen P. Kidd, Dongqing Yang, Gerald J. Atkins

## Abstract

Osteomyelitis associated with periprosthetic joint infection (PJI) is a serious and growing complication of orthopaedic joint replacement surgery, with a failure to cure rate within 2-years of between 15-46%. PJI is a severe chronic infectious disease with a 20% recurrence rate and culture-negativity in 7–39% of cases, of which 14–27% are PCR-positive, findings that remain poorly explained. Osteocytes, the most numerous bone cell type, have been identified as potential long-term intracellular reservoirs for *Staphylococcus aureus*, shielding it from antimicrobial treatment. Here we demonstrate that in chronic intra-osteocytic infections, *S. aureus* consistently adopts a viable-but-non-culturable (VBNC) state, capable of spontaneous host cell escape and reversion to active growth. Whole-genome sequencing identified single nucleotide polymorphisms (SNPs) in gene pairs between clones. Two revertant clones exhibited SNPs in *srrB* and *murA2*, and five exhibited SNPs in *garR* and upstream of *tatB*. While the functional importance of these changes is uncertain, they suggest that diverse genetic mechanisms underly entrance into and exit from the intracellular VBNC state. These findings provide a compelling explanation for culture-negative, PCR-positive osteomyelitis and have direct implications for the diagnosis and treatment of PJI.

## Introduction

Periprosthetic joint infection (PJI) is a serious and growing complication of orthopaedic joint replacement surgery. A major associated problem is the diagnosis of chronic and recurrent infections and the identification of the pathogen, with culture-negative results reported between studies of 7-39% [1–4]. In 14-27% of culture-negative clinical samples, bacteria can be detected at the DNA level using PCR approaches from fluids drained from the joint space [5]. Possible confounding causes for culture-negative infections include poor site and tissue selection for sampling or inadequate processing approaches that do not appropriately disrupt the host tissue, or which damage the pathogens therein [6]. Additionally, biofilm-resident bacteria may not be easily dispersed and some pathogens do not grow on standard media [1, 7]. Beyond these challenges, there is a potential that some bacteria were appropriately sampled but remain undetected as they enter a viable but non-culturable (VBNC) state within tissues that are not readily or currently assayed by PCR. This remains to be conclusively demonstrated under controlled conditions relevant to chronic osteomyelitis. Whilst the presence of the VBNC growth phenotype is known to be involved in implant-associated biofilm-resident bacteria in cases of PJI, this alone cannot explain the current rate of culture-negativity in clinically apparent infections. These infections are often recalcitrant to revision surgery (5.7-16.2% of cases), which can involve either debridement, antibiotic treatment and irrigation with implant retention (DAIR) or the complete removal of the biofilm-laden implant in a staged revision arthroplasty procedure [8–10].

*Staphylococcus aureus*, the most common pathogen in osteomyelitis [11, 12], has been connected to a 4-fold higher treatment failure rate than the next most-common pathogen, *Staphylococcus epidermidis* [13], and has been shown to be capable of forming VBNC in biofilm [14, 15]. It has been previously demonstrated that intra-osteocytic infection of *S. aureus* is a component of PJI-associated osteomyelitis [16–18] and that *S. aureus* adapts to its intracellular environment by forming the quasi-dormant small colony variants (SCV) [16, 19, 20], which have been found in 34% of Staphylococcal PJI cases [13]. Since it is challenging to study intracellular infections *in vivo*, *in vitro* models have been developed to study intracellular *S. aureus* infections in osteocytes, including human primary cells and the Saos-2 cell line [6, 16, 20–25]. Results from a limited number of Staphylococcal osteomyelitis models have indicated that the inflammatory response of osteocyte-like cells is likely dependent on the intracellular bacterial concentration [16, 20]. Furthermore, we recently showed that the loss of culturability post-infection of *S. aureus* in osteocytes did not correspond to a decrease in bacterial burden, as determined by quantification of bacterial DNA load [6, 25].

The VBNC state is believed to be a survival mechanism that bacteria adopt within stressful environments, with phenotypic and potential mechanistic overlap with other growth phenotypes conducive to persistence. One of these, the triggered persister growth phenotype, is defined as a subpopulation that survives extreme stress (usually bactericidal antibiotic treatment or nutrient deprivation) through reversible phenotypic adaptation, and which have an extended lag phase during exposure to and immediately following the removal of the stressor [26]. The VBNC phenotype can similarly be induced by stress although they are distinguished from triggered persisters by the lack of culturability regardless of incubation duration and the fact that growth reactivation is not necessarily triggered promptly or at all by the removal of the inducing stressor [27]. Distinct from the stress-induced, triggered persister phenotype, both spontaneous persisters and VBNC bacteria may exist as subpopulations of a monomicrobial community in the absence of stress. These two growth phenotypes can be differentiated once again by their ability to form colonies and the dynamics of their reversion to a normal growth phenotype. VBNC *S. aureus* are notably capable of replication and metabolic activity whilst remaining non-culturable, although at a lesser rate than the normal growth phenotype [28].

In this study, we observed the bacterial adaptation and infection dynamics, and the host cell innate immune response to chronic infections with clinical isolates of *S. aureus* at different bacterial concentrations in human primary osteocytes *in vitro*. For the results to be consistent with clinical observations, it was hypothesised that the bacterial community would become non-culturable, coupled with an impaired host immune response. Further, non-culturability and reversion to an active growth state would likely be due to one or more autogenous genetic changes, which would be detectable via whole genome sequencing.

## Materials and Methods

### Cell culture

Normal human bone explant-derived cells (NHBC) were cultured from cancellous bone pieces taken from patients undergoing joint replacement surgery and differentiated to osteocyte-like cells, as previously described [16, 29, 30]. Briefly, bone pieces were cultured in growth media, consisting of αMEM (Gibco, Billings, USA) supplemented with 10% v/v foetal calf serum (FCS), 50 mg/ml ascorbate 2-phosphate and standard tissue culture additives (10 mM HEPES, 2 mM L-glutamine, penicillin/streptomycin each 1 unit/ml, all from Thermo-Fisher, Parkville, Australia) at 37°C/5% CO_2_. The cells were either stored in liquid nitrogen or directly passaged to a maximum of 5 passages. For experimentation, cells were seeded at a density of 3×10^4^ cells/cm^2^ in 48 well plates and switched to a differentiation media (growth media, but with 1.8 mM potassium di-hydrogen phosphate (Sigma, St Louis, USA) and only 5% v/v FCS), for 28 days. Cells were maintained with bi-weekly media change and transferred to an antibiotic free media for 24 hours before an infection assay. Each experiment was conducted with at least 2 different donor cells. All the usage and operation procedures for collecting human bone biopsy specimens were approved by the institutional Human Research Ethics Committee.

### Bacterial Culture and intracellular infection assay

All clinical *S. aureus* strains tested were taken from patient’s wound or bone undergoing treatment for DFI and grown in nutrient broth (NB), consisting of 5g/l NaCl, 3 g/l beef extract and 10 g/l peptone (Chem-Supply, Gillman, Australia), on a 37°C/200 rpm shaking platform. The bacterial viable cell concentration was estimated from a colony forming unit (CFU)/ ml vs OD_630_ nm standard curve, then validated by plating dilutions on NB agar, consisting of NB with 1.5% w/v bacteriological agar (Sigma-Aldrich, St Louis, USA), As previously described [20], bacteria were pelleted at 10,000 × g for 10 min, then resuspended in sterile PBS to the target density for the various multiplicities of infection (MOI) according to the requirements of specific experiment, as determined by a spectrophotometric standard curve. The cell media was replaced with the bacterial inoculum in PBS for 2 h in a humidified incubator in an atmosphere of 5% CO_2_, 21% O_2_ at 37°C. Afterwards, cells were washed three times with sterile PBS and incubated with antibiotic free media containing 10 g/l lysostaphin (Sigma-Aldrich). Culture supernatants were verified to be sterile by agar plating at 24 h post-infection. Media were changed every 7 days during the experimental periods. During the second feed (15 DPI) the medium was changed to antimicrobial-free differentiation media (lysostaphin not added). Cultures were monitored daily for evidence of extracellular growth. Supernatants from wells yielding bacterial re-growth were plated onto NA and stored as revertant clones for DNA extraction and sequencing analysis, as described below.

### DNA and RNA processing

For the purposes of real-time and droplet digital PCR analysis, DNA was extracted with the Viagen DirectPCR® DNA Extraction System^TM^ and 1% proteinase K (Australian Biosearch, Wangara, Australia) for 10 min at room temperature, 30 min at 55°C and 15 min at 85°C. RNA was isolated using TRI reagent (Life Technologies, Grand Island, USA) and complementary DNA (cDNA) templates were prepared using the iScript RT kit (BioRad, Hercules, USA), as per manufacturer’s instructions. Both were extracted in biological quadruplicates.

### Real time-PCR and digital droplet PCR

Real-time-PCR reactions were performed using Forget-Me-Not™ EvaGreen® qPCR Master Mix (Biotium, Fremont, USA) on a CFX Connect Real Time PCR System (BioRad) in technical replicates. The sequences of the oligonucleotide primer sets targeting each gene are listed in **Table S1**, Probe sequence for *sigB* used was: 5’-[FAM]caXgtgXgcgtatcggtttaagtcaaatgc[BHQ1]-3’’. Gene expression data was normalised to the geometric mean of *ACTB* and *HPRT1* using the 2^−ΔCt^ method. All primers were designed in-house and purchased from Sigma-Aldrich.

Digital droplet PCR was performed essentially as described [6]. In brief, DNA was digested with Ncol-HF™ (New England BioLabs, Notting Hill, Australia). Reactions were prepared with ddPCR Supermix for Probes (No dUTP) and Droplet Generation Oil for Probes as per manufacturer’s instructions with the QX200 Droplet Digital Generator, C1000 Touch Thermocycler and QX200 Droplet Reader (all BioRad, Hercules, USA).

### Measurement of host cell viability via live/dead staining

As described previously [24], cells were cultured on Cell Imaging Plates (Eppendorf, Hamburg, Germany) as described above. After 1-, 7- and 21-days post infection, the cells were incubated for 20 min with Ethidium Homodimer III (Biotium) and eBioscience^TM^ Calcein Violet 450 AM Viability Dye. Confocal images were taken with an Olympus FV3000 confocal microscope (Olympus, Tokyo, Japan) and analysed with Fiji ImageJ to obtain the relative intensity.

### Quantification of culturable intracellular bacteria numbers

At 1-, 7- and 21-days post infection, 4 wells were lysed in sterile water for 20 min at 37°C and 1:10 serial diluted. Lysates and dilutions were plated on NB agar and incubated up to 14 days at 37°C in a 5% CO_2_ humidified incubator and examined after 24 h and before disposal at least 14 days of incubation to ensure capture of culturable bacteria with extended lag phases such as SCVs.

### Immunostaining

Cells were cultured on Cell Imaging Plates (Eppendorf) as described above. After 21 days of infection, cells were fixed in Theralin tissue fixative (Grace Bio-Labs, Bend, USA) at room temperature for 3 days and stained, as described previously [16]. Briefly, cells were stained using a 1:100 dilution of *S. aureus* targeting antibody (AB37644; Abcam Inc., Cambridge, USA) at 4°C for 1 h to detect the pathogen. Primary antibody signals were tagged by Alexa Fluor 647-conjugated rabbit anti-mouse IgG (H+L) secondary antibody (A-21239; Life Technologies, Inc.) for visualization. Using matched-isotype control antibodies (14-4742-81; ThermoFisher) for primary detection, the specificity of antibody staining was validated by following the same staining procedures. DAPI was used to counterstain nuclei. All samples were imaged with an Olympus FV3000 confocal microscope (Olympus).

### Transmission Electron Microscopy

Individual wells from two donors infected with either 142B or 22W *S. aureus* or uninfected controls from 1-, 7- and 21-days post infection were fixed in 1.25% v/v glutaraldehyde, 4% w/v sucrose and 4% w/v paraformaldehyde and PBS (pH 7.2), and then processed in 2%, wt/vol, osmium tetroxide in water. Cell pellets were gradually dehydrated in EtOH, embedded in resin blocks, and 5-µm sections were taken for transmission electron microscopy (TEM; Philips CM200) imaging as previously described [16, 20].

### RNA Sequencing

Total RNA was isolated (as above) from human primary osteocyte-like cultures generated as described above, 27 days post-infection with *S. aureus* WCH-SK2 (original MOI 40) at a stage where no extracellular bacterial growth had been observed for 14 days. RNA-Seq was performed by the Australian Genome Research Facility (AGRS, Melbourne, Australia) using TruSeq Stranded mRNA (Illumina) sample preparation with sequencing using a NovaSeq 6000 sequencing system (Illumina), with at least 20 million 100 bp single reads per sample. Bioinformatics included quality and adapter trimming, alignment, quantification and normalisation (AGRS).

### Whole Genome Sequencing

Bacteria were cultured on Tryptone Soya Agar (Oxoid) solid medium and incubated at 37°C for 16-20 hours. Single colonies were used to inoculate Terrific broth (Invitrogen) and cultured overnight at 37°C at 200rpm. Seed cultures were back diluted to an OD_600_ of 0.05 in Terrific broth and grown until early mid-log. Bacteria were then concentrated and washed in 1x PBS. *S. aureus* genomic DNA (gDNA) was extracted using the Qiagen Genomic-tip 20/G kit in combination with the genomic buffers set (Qiagen). The lysis process was performed as per the manufacturer’s instructions (Qiagen) with the addition of Lysostaphin. Resultant gDNA was quantified by Nanodrop (Thermofisher) and Qubit fluorometer (Invitrogen). Fragment length and DNA integrity were assessed by a 4150 TapeStation system (Agilent). Library preparation was performed using Oxford Nanopore Technologies Rapid Sequencing DNA V14 barcoding kit (SQK-RBK114-24, Oxford Nanopore), as per the manufacturer’s instructions. Sequencing was performed on a MinION sequencer with an R10.4.1 chemistry flow cell (Oxford Nanopore). Post-sequencing, POD5 files were re-base-called with Oxford Nanopore’s Basecaller, Dorado (dna_r10.4.1_e8.2_400bps_sup@v5.0.0) using the Super accurate (SUP) model.

Genome assembly was performed using Hybracter (version 0.10.0), a long read bacterial genome assembly pipeline [31]. The Hybracter-long workflow was used, with minimum length and minimum quality options set to 1000 base pairs and 15 respectively. The resultant fasta files were annotated using Bakta (version 1.9.1) and its pre-configured database [32].Variant calling was performed using Oxford Nanopore Technology’s haploid variant caller, Medaka (version 2.0.1), using the WCH-SK3 parental Hybracter assembly as a reference. Resultant .vcf files were mapped to gene names using Bedtools intersect. Aligned reads at all variant positions identified by Medaka were visually inspected to identify artefactual variants using the Integrative Genomics Viewer (IGV) [33]. All artefactual variants had quality scores <30 and all confirmed variants had quality scores >30. We report only confirmed variants.

### Statistical analysis

Each experiment has been conducted with a minimum of three biological and two technical replicates. Two-way ANOVA with Tukey’s post-hoc tests were used. All analysis was performed using GraphPad Prism software (GraphPad Software, La Jolla, USA) Values for p < 0.05 were considered significant.

## Results and Discussion

Across a 28-day time course of infection with high multiplicities of infection (MOI) of either of two clinical isolates of *S. aureus*, colony forming unit (CFU) counts decreased, with no culture being detectable by 21 days post-infection (DPI; **Fig. 1A**). However, intact bacteria were still detectable by TEM (**Fig. 1B**) and immunofluorescence imaging (**Fig. 1C**), indicating that bacteria remained present. The structure of intracellular bacteria under TEM had varied morphology, some being distinct from the spherical shape of extracellular bacteria in a normal growth condition, perhaps indicating variation of cell wall rigidity or integrity underlying the lack of culturability. No such structures were seen in uninfected cells. Despite the altered cellular shapes, the apparent electron density of intracellular bacteria and basic cell wall morphology were consistent with the extracellular bacteria in positive control samples (co-cultures where bacteria grew into the media before being fixed for imaging). Such irregular bacterial shapes have previously been described in VBNC *S. aureus* [28], as well as during *S. aureus* invasion of the osteocyte lacunocanalicular system [34].

**Figure 1:**
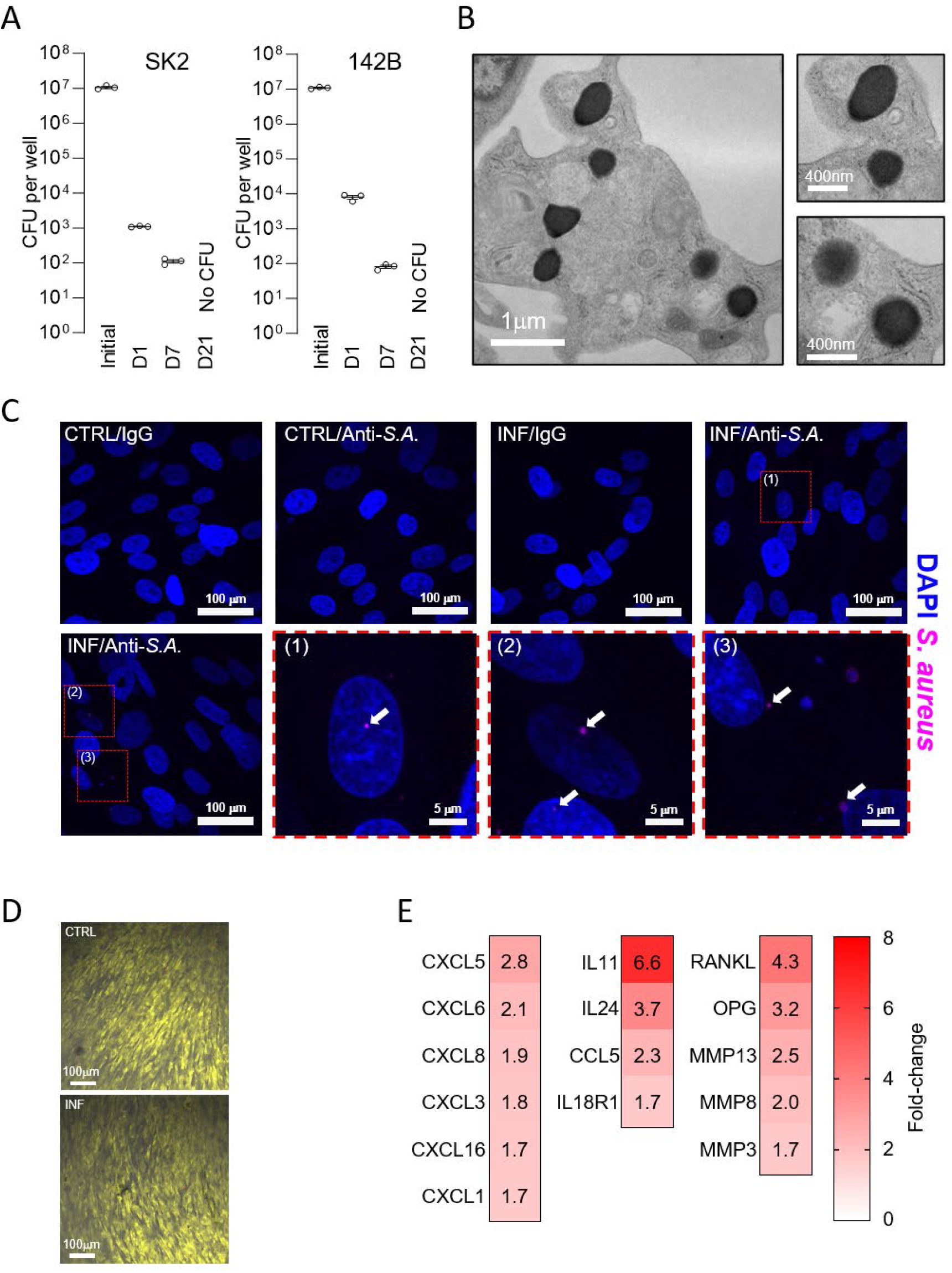
Characterisation of the dynamics of high dose *in vitro S. aureus* infection of cultured human primary osteocyte-like cells. (A) Time course of colony forming units (CFU) cultured either from the inoculum (initial) or primary osteocyte lysates (1-, 7- and 21-days post-infection (DPI)) of two clinical isolates (WCH-SK2 and 142B). (B) Transmission electron microscopy (TEM) images of 142B *S. aureus* infected primary human osteocytes at 21 DPI, showing intracellularly resident *S. aureus.* (C) Immunofluorescent confocal microscopy images of either WCH-SK2 infected (INF) or uninfected (CTRL) cultured human primary osteocytes. Anti-*S. aureus* (Anti-SA) or an isotype control immunoglobulin G (IgG) were used as primary antibodies (*pink*). 4’,6-diamidino-2-phenylindole, (DAPI, *blue*) was used as a nuclear counterstain. Three inset images are indicated by dashed boxes. (D) Fluorescent confocal images demonstrating host cell viability 21 DPI with Calcein 450 AM Viability Dye (live cells, *yellow*) and Ethidium homodimer III (dead cells, *red*). (E) Heat plot showing relative mRNA expression levels of a selected range of immunomodulatory CXCL and CCL chemokines, cytokines, cytokine receptors, bone remodelling genes and matrix metalloproteinases, between MOI 40 WCH-SK2 *S. aureus*-infected human primary osteocytes at 27 DPI and uninfected control cultures.

Host cell viability was not impacted by long-term infection (**Fig. 1D**), confirming that the observed decline in culturable bacterial burden was not due to the death of host cells eliminating a niche of persistence but instead the survival of a resident subpopulation of non-dividing bacteria. This is consistent with our previous observations using an osteocyte-like cell line infected with *S. aureus*, with no impact on cell viability following a low dose of infection [20]. This observation is also consistent with our previous work indicating that chronic recurrent osteomyelitis in PJI can be facilitated through the long-term persistent infection of viable osteocytes [16].

Indicative of an ongoing host cell response to chronic infection, RNA-Seq analysis of the host cell responses after 4-weeks of infection showed an upregulation of the CC chemokine ligand (CCL) genes *CCL3*, *CCL5* and *CCL8*, the CXC chemokine ligand (CXCL) genes *CXCL2*, *CXCL3*, *CXCL5*, *CXCL6* and *CXCL16,* and the interleukin (IL) genes *IL1A*, *IL15* and *IL6* (**Fig. 1E**). These findings are reminiscent of those in an acute infection model of human osteocytes and of bone infected *ex vivo* [16], although both the relative array and strength of expression of immune regulators were reduced in the current chronic model. The chemokine/cytokine response pattern here would be expected to mainly attract and activate neutrophils, which play an important role in immune defence but also cause damage to the infected tissue by the release of, amongst other mechanisms, reactive oxygen and reactive nitrogen species. Additionally, some of the induced immune mediators are known to modulate bone remodelling. While CXCL16 and IL6 promote osteoblast differentiation and migration [35, 36], CXCL2 inhibits osteoblast differentiation [37], while IL-15 is reported to increase osteoblast and osteoclast apoptosis [38]. CCL5, CXCL2, IL-1α and IL-6, on the other hand, promote osteoclastogenesis and bone resorption [39–42], with IL-1α additionally inhibiting collagen synthesis [43]. Collectively, the observed gene expression response profile indicates a shift towards bone catabolism. Indeed, bone matrix collagen has been found to be degraded in bone from PJI patients, which was linked to an increased expression of collagenases, such as matrix metallopeptidase (MMP)-13 and MMP1 in both PJI patient bone, as well as in osteocytes exposed acutely to *S. aureus* [44].

The change in culturability was found to be broadly similar between high and low dose bacterial infections, with all infections ultimately resulting in no culturable bacteria within host cell lysates at the final time point (**Fig. 1A, 2A-B**). The exception to this was low dose infection with a clinical osteomyelitis strain designated 142B, where there was an increase in retrieved CFU between 1 and 7 DPI, which then conformed to the otherwise observed pattern by 21 DPI (**Fig. 2B**). In contrast to CFU numbers, total bacterial DNA levels quantified by droplet digital PCR (ddPCR) plateaued after an initial decrease and were still measurable at the same point that no CFU were recovered (**Fig. 2A**). Furthermore, bacterial numbers quantified via ddPCR were always higher than the CFU count, and in one case were even in excess of the inoculum dose, demonstrating intracellular replication. This indicates that a substantial portion of the intracellular bacterial community had reached a non-culturable state even by 1 DPI, as the number of bacterial genomes per well at this time-point was orders of magnitudes higher than the CFU numbers recovered. While not all *S. aureus* DNA measured was necessarily from VBNC, the combination of persistently high copy numbers of DNA across multiple weeks, the presence of cellular structures resembling intracellular bacteria visible under TEM and fluorescent *S. aureus* surface immunostaining by 21 DPI (**Fig. 1B-C**), support the presence of a persistent and VBNC intracellular bacterial community. The decrease in intracellular bacterial DNA levels over time in concert with consistent low levels of culture from lysates suggest the presence of a subpopulation of the initial intracellular population capable of successfully adapting to this environment for long-term persistence. Interestingly, a similar residual ddPCR count was observed for both strains tested, perhaps indicative of a final population size of persisters that can be maintained within this host cell type and number. Some bacterial strain variation was evident, as the 142B infection was seen to reach parity between genome copy and CFU counts at 7 DPI, although both strains transitioned to a completely non-culturable persistent population by 21 DPI. Further, this transient parity also strongly indicates that the DNA measurements were from predominantly viable cells. Finally, it is significant that this phenomenon of bacterial transition into a VBNC state with a steady bacterial number, mediated by intra-osteocytic residence, is robust and not strain-specific. All 11 *S. aureus* strains were tested under the same conditions and showed a CFU reduction of between 1.98-3.11 logs after 7 days and became non-culturable after 14 days of infection, while bacterial mRNA only dropped 0.03-1.31 logs after 7 days and was still measurable in all strains (0.35-1.69 log-reduction) after 14 days; the mRNA reduction was always less compared to the decrease in CFU numbers (**Suppl. Fig. 1**).

**Figure 2:**
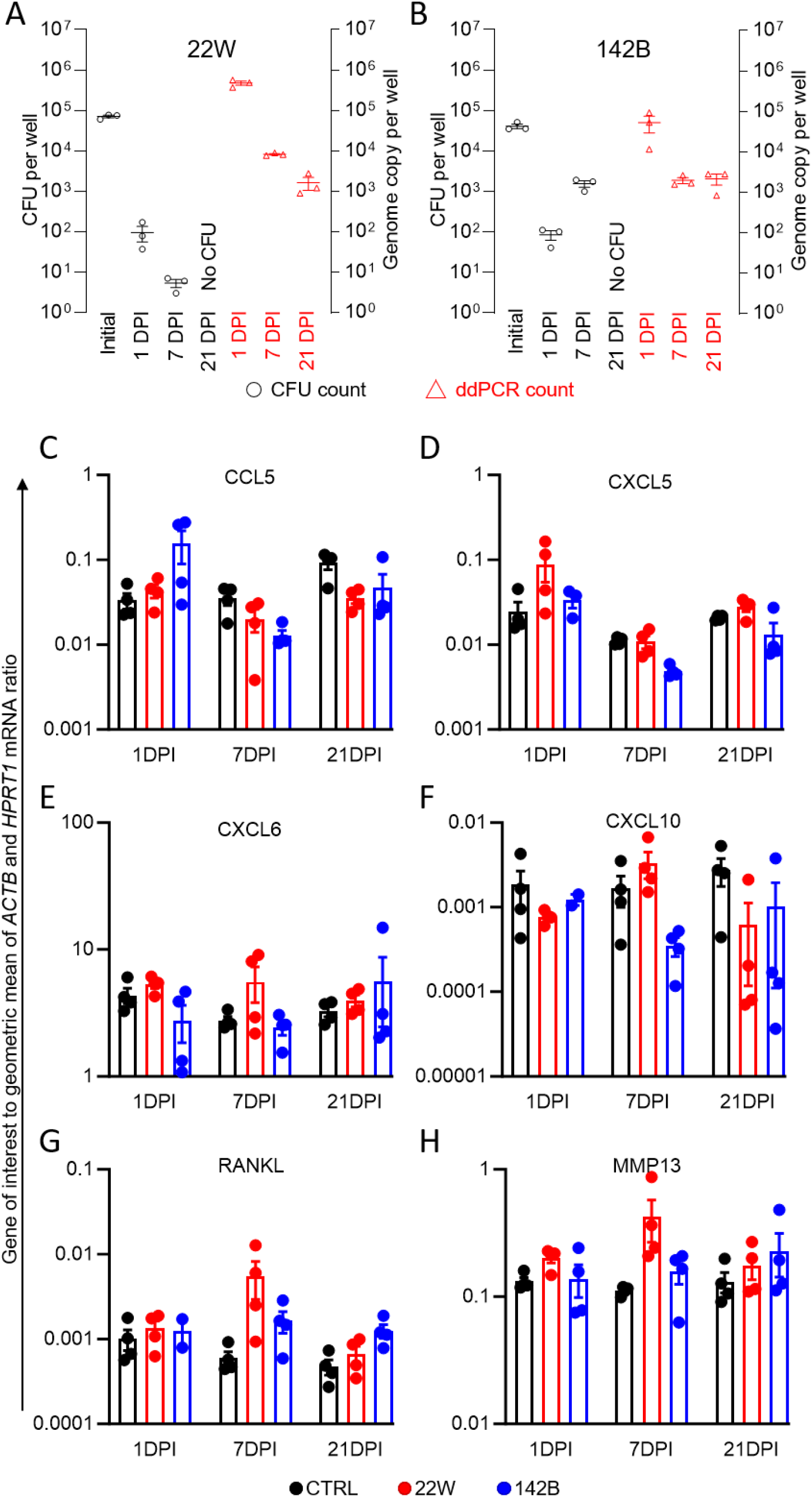
Characterisation of bacterial growth dynamics of low dose *in vitro S. aureus* infection of cultured human primary osteocytes. (A) Time course of colony forming units (CFU, *black*, left axis) cultured either from the inoculum or osteocyte lysates (1-, 7- and 21-days post-infection (DPI)) of two clinical isolates (22W and 142B), and corresponding numbers of the bacterial single copy gene *sigB* as an indication of bacterial genome copy number from host cell lysates were quantified via droplet digital polymerase chain reaction (ddPCR, *red*, right axis). (C-H) Relative mRNA expression levels normalised to the geometric mean of *ACTB* and *HPRT1* of immunomodulatory chemokines *CCL5*, *CXCL5*, *CXCL6* and *CXCL10* (C-F), the osteoclastogenic factor *RANKL* (G) and the collagenase *MMP13* (H).

While host-cells infected with a relatively high bacterial dosage showed an innate immune response that was significantly different from uninfected cells across the entire time course studied herein (see **Fig. 1E**), those with a lower bacterial burden did not show a significant response in any gene examined (**Fig. 2C-H**). A possible factor for this difference could be the low MOI for the infection used, as has been previously demonstrated in the osteocyte-like Saos-2 cell line, with orders of magnitude higher doses of *S. aureus* (strain WCH-SK2) maintaining a significantly distinct pro-inflammatory response at 15 DPI [20]. Thus, a larger inoculum results in a larger persister community and beyond some threshold of relative abundance this population is incapable of suppressing or evading the host cell immune response. At the low MOI used it would inevitably be the case that not all host cells would be infected and the decline in bacterial DNA across the infection time course supports this, as does the rarity of infected cells observable within TEM sections. The fact that a community of osteocytes harbouring rare, infected cells showed no overall difference in the expression of inflammatory cytokines compared to a similarly sized population of uninfected cells, which were otherwise upregulated in high dose infections, speaks to the failure of a low number of infected cells to elicit a significant paracrine inflammatory cascade. This could cause a failure to recruit towards the site of infection, consistent with the immunologically silent chronicity often seen clinically. The ability to suppress the host cell initiation of an inflammatory response is a key mechanism of long-term intracellular persistence [45]. Together with the relative inaccessibility afforded by the anatomical niche of deep bone osteocytes to the cell-mediated arm of the immune system, this perhaps helps explain why *S. aureus* is one of the most common causative organisms for osteomyelitis.

We have previously demonstrated *in vitro* the ability for these resident quiescent bacteria within the osteocyte niche to reactivate to a culturable state spontaneously [23] and survive clinically relevant antimicrobial therapy [25] in a permissive environment, further suggestive of the importance of this pathway in chronic, relapsing PJI. In this study, culture of *S. aureus* beyond 3 weeks also resulted in reactivation of quiescently-infected wells. To investigate this phenomenon, colonies from spontaneously reverted chronic non-culturable infections within osteocytes (at 24 DPI) were analysed by WGS and compared to the corresponding parental strain. Revertant clones exhibited one or more single nucleotide polymorphisms (SNPs) that may have caused the phenotypic shifts in growth state (**Table 1**). Two clones exhibited a SNP (T>A) resulting in the amino acid change K161I in the protein product of *srrB*, a gene involved in global virulence factor expression [51] and hypoxia / nitric oxide stress resistance [52]. Previous research indicated *srrAB* mutations were capable of reverting SCVs into a normal growth phenotype [53], such that both genetic changes observed independently support a mechanistic overlap between the SCV and VBNC growth phenotypes. These two clones also exhibited a SNP in *murA2*, encoding UDP-N-acetylglucosamine 1-carboxyvinyltransferase essential for cell wall peptidoglycan synthesis [54]. Five additional clones exhibited a SNP in *graR*, a gene associated with resistance to vancomycin [55]. These same clones exhibited a SNP in an upstream region of *tatB*, encoding a component of the twin-arginine protein secretion pathway TatA/TatB, the folded protein transmembrane translocation system [56]. It remains to be seen whether one or more of the SNPs identified were mechanistically responsible for either entry or exit into the intracellular VBNC state, and whether others exist. We propose that the existence of multiple mechanisms is highly likely, given the versatility and success of *S. aureus* as a human pathogen and the diversity between clones observed here; however, these genetic changes are the first described in the context of intracellular VBNC and osteomyelitis.

**Table 1:** SNPs identified in WCH-SK3 revertant clones.

| *Variant Position* | *Isolates with variant* | *Nucleotide change* | *Amino acid change* | *Gene* | *Gene Function* |
| --- | --- | --- | --- | --- | --- |
| 1475136 | SK3-R1<br>SK3-R2 | T > A | K161I | *srrB* | Two-component system sensor histidine kinase involved in global virulence factor regulation.[51] |
| 2093513 | SK3-R1<br>SK3-R2 | A > G | M144T | *murA2* | UDP-N-acetylglucosamine 1-carboxyvinyltransferase essential for cell wall peptidoglycan synthesis.[54] |
| 340587 | SK3-R3<br>SK3-R4<br>SK3-R5<br>SK3-R6<br>SK3-R7 | G > A | - | Upstream of<br>*tatB* | Twin-arginine protein secretion pathway components TatA and TatB, the Folded protein transmembrane translocation system.[56] |
| 658487 | SK3-R3<br>SK3-R4<br>SK3-R5<br>SK3-R6<br>SK3-R7 | T > G | F59V | *graR* | Response regulator transcription factor graR/apsR; mutations associated with antimicrobial resistance, e.g. vancomycin resistance.[55] |

## Funding Statement

This work was supported by an Ideas Grant (ID 2011042) from the National Health and Medical Research Council of Australia (NHMRC). NJG was supported by an Australian Postgraduate Award. ARZ and MAH were supported by University of Adelaide Postgraduate Awards.

---

**Supplementary Figure 1:**
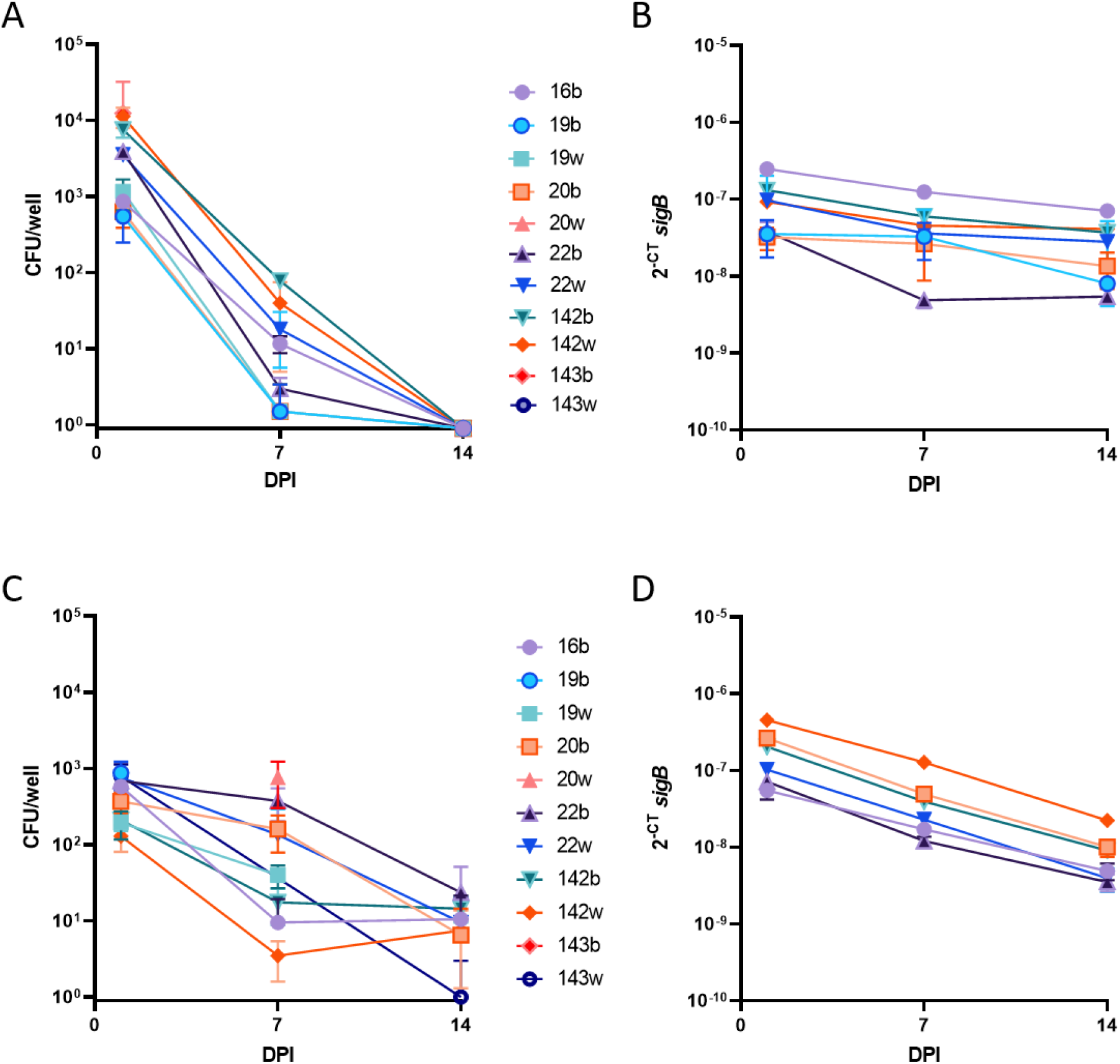
Intracellular persistence characteristics of 11 *S. aureus* strains in human osteocytes *in vitro.* Human primary cell-derived osteocytes (A/B) or Sa0S2-0Y cells (CID) were infected with the various strains isolated from diabetic foot infection samples, adjusted for MOI = 1, as described in Materials and Methods. Cells were harvested at the times indicated and plated for intracellular CFU or prepared for total RNA isolation and RTPCR for bacterial *sigB* expression. Data shown are mean values of biological quadruplicates. DPI = days post-infection. CFU = colony forming units.

**Supplementary Table 1:**
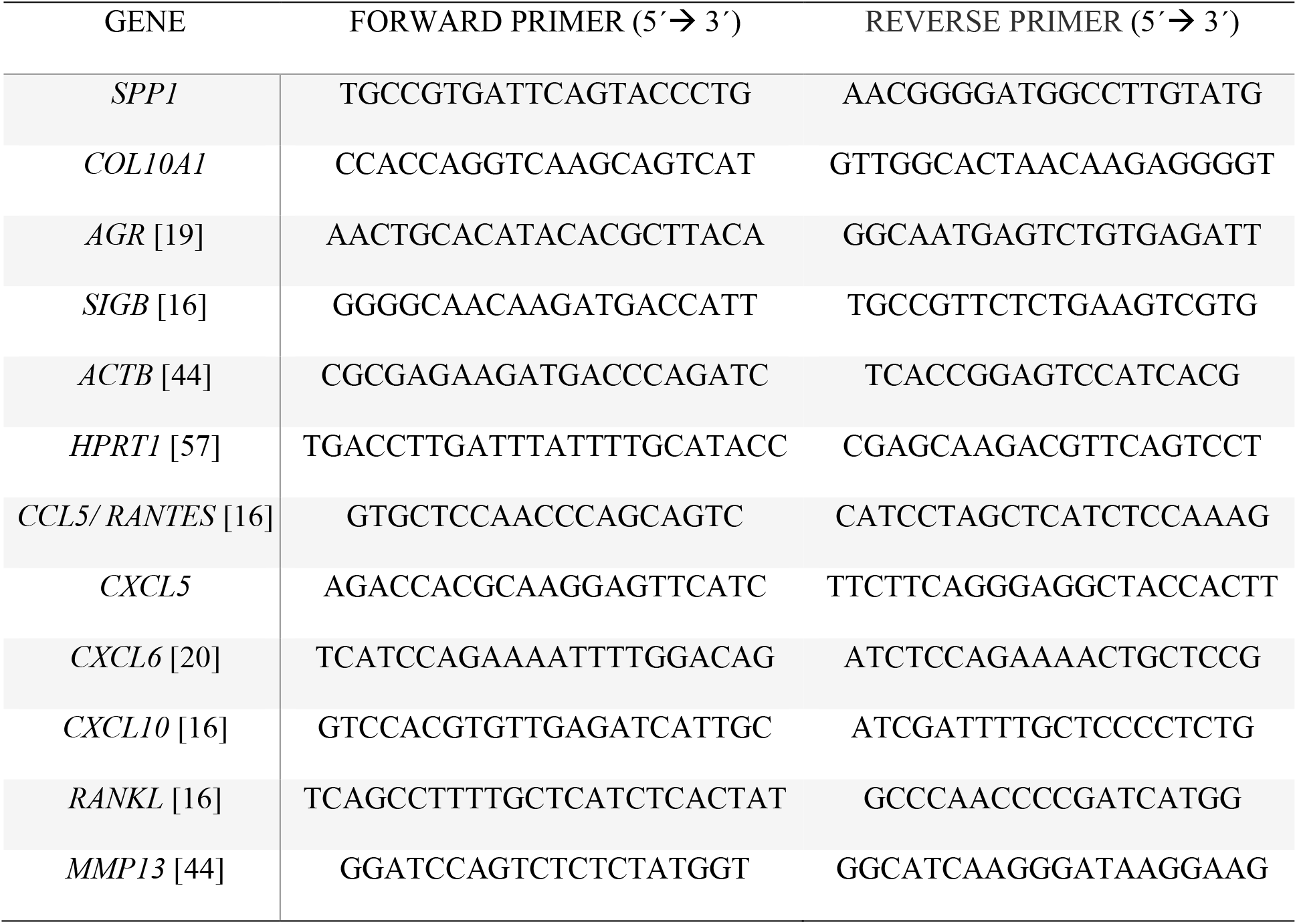
Oligonucleotide primer sequences used for qPCR and ddPCR.

## Notes

### Competing Interest Statement

The authors have declared no competing interest.

### Summary of Updates

In this version, we have substantially reworked the paper to improve clarity and expression. Importantly, we have updated the data pertaining to whole genome sequening of S. aureus revertant clones. Having previously only performed the assay on one clone (i.e. n=1), we sought to ensure there was reproducibility in the outcomes and provide data from biological replicates. The new genetic events identified from n=7 revertant clones (see new Table 1) we believe reflect a convergent evolutionary outcome related to an intracellular VBNC state.

